# Hydrodynamic trapping measures the interaction between membrane-associated molecules

**DOI:** 10.1101/357277

**Authors:** Victoria Junghans, Jana Hladilkova, Ana Mafalda Santos, Mikael Lund, Simon J. Davis, Peter Jönsson

## Abstract

How membrane proteins distribute and behave on the surface of cells is determined by the molecules’ interaction potential. However, measuring this potential, and how it varies with protein-to-protein distance, has been challenging. We here present how a method we call hydrodynamic trapping can achieve this. Our method uses the focused liquid flow from a micropipette to locally accumulate molecules protruding from a lipid membrane. The interaction potential, as well as information about the dimensions of the studied molecule, are obtained by relating the degree of accumulation to the strength of the trap. We have used this to study four representative proteins, with different height-to-width ratios and protein properties; from the globular streptavidin, to the rod-like immune cell proteins CD2, CD4 and CD45. The obtained data illustrates how protein shape, glycosylation and flexibility influence the behaviour of membrane proteins as well as underline the general applicability of the method.

## Introduction

Both the interaction and the transfer of information between cells is regulated via membrane proteins acting among others as receptors, adhesion molecules, or ion transporters. Vital for this is the interaction with other membrane proteins. These interactions can be either permanent or transient, allowing for changes in the oligomeric state of the molecules.^1^ However, little is known about the physicochemical forces acting between membrane-anchored molecules and to date it has been problematic to study these interactions between molecules bound to lipid bilayers. We recently showed that the liquid flow through a micropipette can be used to trap and accumulate molecules in a supported lipid bilayer (SLB), a technique we call hydrodynamic trapping.^2,3^ The flow acts on molecules protruding from the lipid bilayer with a drag force whose magnitude can be accurately controlled by varying the flow rate through the pipette and distance to the SLB. It is in this way possible to change the local concentration of membrane-bound molecules by orders of magnitude.^2^ We here show how hydrodynamic trapping can be used to measure the interaction potential between membrane-anchored molecules, which combined with Metropolis-Hastings Monte Carlo (MC) simulations allow us to draw a series of general conclusions of how protein shape, glycosylation and flexibility affect the interaction between membrane-associated molecules.

The basis of the method is the following. A micropipette is positioned above an SLB containing one or two sorts of proteins at an average surface concentration of 100 to 1000 molecules/µm^2^ (Figure 1A). Negative pressure is applied through the micropipette and the proteins start to accumulate (Figure 1B, C). The higher the applied pressure, and/or the closer the micropipette is to the surface, the higher is the force acting on the molecules resulting in an increased protein accumulation (Figure 1D). By relating the accumulation to the strength of the hydrodynamic trap we can estimate: (i) the approximate dimensions of the molecule, and (ii) the intermolecular forces between the molecules and how this varies with protein-to-protein distance.

**Figure 1.**
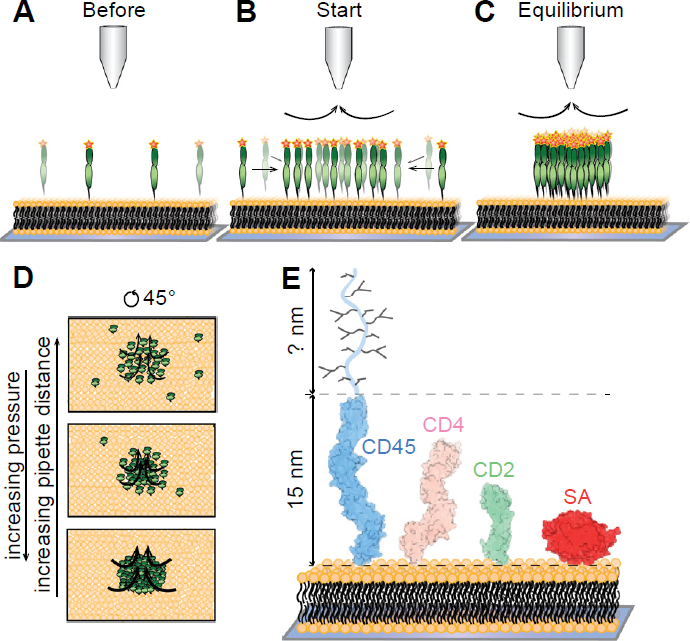
Schematic illustration of the experiments and the studied molecules. **A-C)** Hydrodynamic trapping of membrane-anchored molecules in an SLB is achieved by applying negative pressure through a micropipette (tip radius ~1 μm). This results in accumulation of the molecules in the SLB. Not drawn to scale. **D)** The accumulation depends on the applied pressure and the distance between the pipette and the SLB. **E)** Dimensions of the studied molecules: human CD45 (CD45d1-d4: PDB 5FMV + mucin-like region), human CD4 (PDB 3T0E), rat CD2 (PDB 1HNG) and SA (PDB 3RY2).

We selected four membrane-anchored molecules of varying shape, level of glycosylation and flexibility for our method to investigate. These molecules were the B vitamin biotin-binding protein streptavidin (SA), and the immune-cell membrane proteins CD2, CD4 and CD45RABC (CD45). SA has a globular shape (width and height of ~5 nm)^4^ and is used in numerous biotechnological and diagnostic applications.^5^ The three immune-cell proteins have, presumably, rod-like shapes, with a cross-sectional width of ~3 nm, but vary in height from approximately 7 nm to potentially over 40 nm, see Figure 1E.^6–9^ CD2 is the smallest of these proteins (height ~7.5 nm),^6^ but is also heavily glycosylated, and is important for stable cell-cell adhesion between immune cells. CD4 (height ~11 nm) is known to interact with MHC class II molecules on antigen presenting cells (APCs) but with a very low affinity,^8,10^ heightening the T-cells sensitivity to foreign antigen. CD45 is a receptor type tyrosine phosphatase regulating T-cell signalling, suggested to be excluded from the close contact area between immune cells due to its large protein size and heavy glycosylation (height >15 nm).^7,11^ The orientation and interaction between these molecules are not only interesting from a model point of view but is also important for how these molecules will distribute and behave in the contact between immune cells, which are key factors to better understand immune-cell signalling.

## Results

### From trapping to interaction curves

A description of the trapping method and how to obtain the interaction potential is described below for the protein SA. A micropipette was positioned a fixed distance above an SLB containing anchored SA, after which a negative pressure was applied through the pipette creating an inward flow and accumulation of SA (Figure 2A and Movie S1). A higher pressure applied through the pipette led to greater accumulation. The relative intensity increase compared to before trapping is shown in Figure 2B, where the intensity is radially averaged around the centre of the trap. This value can in turn be converted to a concentration of proteins at different positions in the trap (see “*Methods - Conversion between fluorescence intensity and protein density*”). Whereas SA has been observed to crystalize at higher surface densities,^12^ this was not the case in the current trapping experiments. However, it was observed that photobleaching trapped SA molecules, under high salt and low pH, resulted in the proteins becoming immobile (Figure S1 and Movie S2).

**Figure 2.**
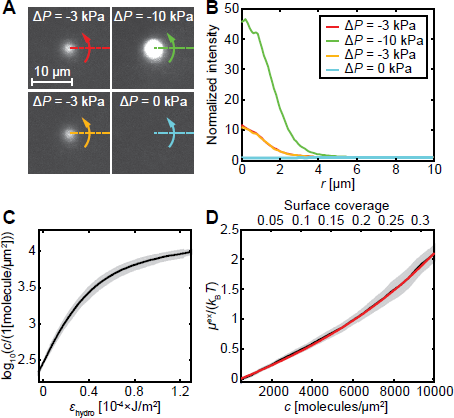
From trapping data to interaction curves. **A)** Images of fluorescently-labelled SA being accumulated on an SLB at different applied pressures, Δ*p*. **B)** Radially-averaged intensity profiles of the data in A. **C)** The interaction curve of SA (mean ± SD). **D)** The experimentally determined excess chemical potential for SA (black; mean ± SEM). The red line is the theoretical curve for a hard disk with a radius of 3.2 nm.

To relate the accumulation of SA to the magnitude of the liquid flow from the pipette we note that the concentration of molecules, *c*, in the trap is at steady state related to the force on a molecule from the liquid flow, *F*_hydro_, by:^2^

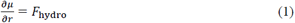

where *μ* is the chemical potential of the molecule at the distance *r* from the centre of the trap. The hydrodynamic force depends linearly on the shear force *σ*_hydro_ according to:^13^

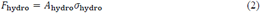

where *A*_hydro_ is the effective hydrodynamic area of the macromolecule. The quantity *A*_hydro_ depends on the size and geometry of the studied molecule, as well as on the surface concentration, *c*, of molecules. Taller molecules generally have larger *A*_hydro_ than shorter molecules, and *A*_hydro_ decreases as *c* increases due to molecules shielding each other from the flow.^13^ The shear force can in turn be determined from finite element simulations (see “*Methods - Determining *ε*_hydro_ and *σ*_hydro_ from finite element simulations*”), and depends on the dimensions of the pipette, the applied pressure through the pipette and the distance between the pipette and the surface. Changing the applied pressure therefore enables the trapping strength to be varied and with this also the number of accumulated molecules (see Figures 2A, B). Combining Eqs. 1 and 2 gives:

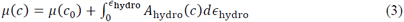

where

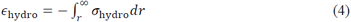

The intermolecular interactions between hard disks of radius *a* are thermodynamically given by the chemical potential, *μ*, of the molecules according to:

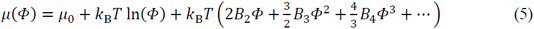

where *Φ* = *c*×π*a*^2^ is the surface coverage at the molecular concentration *c*, *μ*0 is the chemical potential at the standard state and *B*i is the i^th^ virial coefficient of the expansion.^14^ For hard disks in two dimensions the first three virial coefficients take on the following values: *B*2 = 2, *B*3 = 3.128 and *B*4 = 4.258.^14^ Equation 5 will at low coverage approach *μ*(*c*) = *μ*(*c*0) + *k*B*T* ln(*c/c*0), where *c*0 is the concentration before trapping. Inserted into Eq. 3, and setting *A*_hydro_ = *A*_hydro_(0), results in:

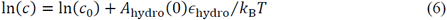

A plot of ln(*c*) vs *ε*_hydro_ (henceforth called an “interaction curve”; Figure 2C), will therefor at low coverage have the slope *A*_hydro_(0)/*k*B*T*. We have previously shown that *A*_hydro_(0) can be estimated using the following empirical expression:^13^

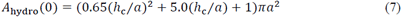

where *h*c is the height and *a* the radius of the molecule. The interaction curve is a characteristic function for each studied protein, which in general is independent of how the curve is obtained (such as the magnitude of the liquid flow, the size of the pipette etc., see also Figure S2).

### The interaction between globular proteins - streptavidin

The initial slope of the interaction curve for SA gave *A*_hydro_(0) = 300±29 nm^2^ (mean ± SD). This value is similar to the theoretical value of *A*_hydro_(0) = 296 nm^2^, obtained using Eq. 7 when modelling SA as a disk with a radius of *a* = 2.8 nm and height *h*c = 5 nm from the crystal structure.^4^ The value for *A*_hydro_ at different concentrations will in addition to *A*_hydro_(0) also depend on the quotient *h*c/*a*.^13^ However, the dependence on *h*c/*a* is weak in the range *h*c/*a* = 1 to 10 (Figure S3) and the function *A*_hydro_(*c*) can therefore be estimated from *A*_hydro_(0). This, together with the values from the interaction curve, can be inserted into Eq. 3 to give the chemical potential *μ*. Figure 2D shows SA´s excess chemical potential defined as: *μ*^ex^ = *μ* – *k*B*T* ln(*c*/*c*0). The red line is a fit of the excess chemical potential to a hard disk model (Eq. 5) yielding a disk radius of 3.2±0.4 nm. This is 0.4 nm larger compared to the crystal structure for SA. The increase in radius is explained by the extra area taken up by the fluorescent Alexa Fluor^®^ 647 groups (on average three per SA molecule), which would correspond to an effective increase in cross-sectional area of the order of 30% in total (Figure S4).

### The orientation and height of CD2, CD4 and CD45 on the lipid bilayer

We next turned to investigate the behaviour of the presumably rod-like proteins CD2, CD4 and CD45 anchored to an SLB. Both the orientation of these molecules on the SLB as well as their effective size when including sugars is, in contrast to SA, not well defined. Trapping experiments for each of the proteins were performed as described previously for SA. Figure 3A shows the interaction curves for the three proteins. CD2 has the shallowest slope at low surface coverage with CD4 in the middle and with CD45 having the highest slope. With a higher slope corresponding to a higher *A*_hydro_(0) value, and thus a larger molecule, this is in agreement with CD45 being the largest followed in size by CD4 and CD2. It is of interest to compare the experimental values for *A*_hydro_(0) that are summarized in Table 1 with the values obtained from Eq. 7 using data from structural studies, assuming the proteins are “standing up”. All immune-cell proteins were approximated as a cylinder with a radius of 1.5 nm, neglecting the effect of the sugar groups on the hydrodynamic area (see Table 1 for values). The effective height of the molecules, *hA*_hydro_, is calculated using Eq. S2 and the experimental *A*_hydro_(0) value. The obtained results are in good agreement with CD2 and CD4 standing upright on the SLB with an effective height within 7% of that obtained from the protein structures (Table 1). The slightly larger height obtained for CD2 can be attributed to the extra drag taken up by its four N-glycans, whereas this deviation would be smaller for CD4 which only has two glycosylation sites.^15^ The full length of CD45 on the other hand has previously been estimated to be approximately 40 nm,^9^ which is considerably larger than the effective height of 22 nm obtained from the trapping data. The extracellular domain of CD45 consists of an N-glycosylated region (CD45d1-d4) with a height of approximately 15 nm, followed by a mucin-like region.^7^ One reason for the low effective height obtained by us might be that the mucin-like region takes up considerably less hydrodynamic force than estimated by Eq. 7. Another possibility is that CD45 is significantly tilted at the low coverage where *A*_hydro_(0) is measured. This also agrees with CD45 being more flexible than the other molecules investigated in this work (see below). However, understanding the exact conformation of CD45 is beyond the scope of the present study.

**Figure 3.**
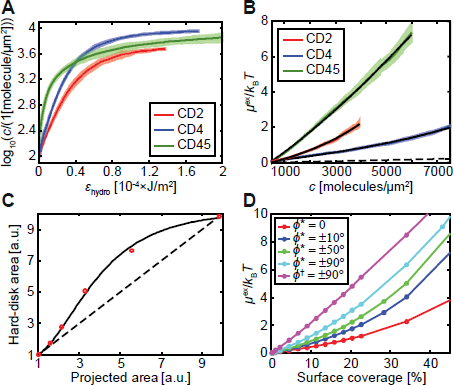
Hydrodynamic trapping of CD2, CD4 and CD45. **A)** Interaction curves for the three proteins (mean ± SD). **B)** The experimentally determined excess chemical potential for the proteins (mean ± SEM). The black lines are fits to the following theoretical models: (i) “hard disks” for CD2 and CD4 and (ii) “±90° rotating rod” for CD45. The dashed line is *μ*^ex^ for a hard disk with a radius of 1.5 nm. **C)** The effective hard-disk area for a glycosylated molecule vs the projected area of the protein + sugars obtained from MC simulations. The dashed line corresponds to the hard-disk area being equal to the projected area and the solid line is a fit to Eq. 15. **D)** Excess chemical potential curves from MC simulations for a rod-like molecule which can rotate an angle *ϕ* around its attachment point. * Molecule consists of three connected spheres. ^†^ Molecule consists of six connected spheres.

**Table 1.** Theoretical and experimental values of the protein radius, *a*, height, *h*_c_, and hydrodynamic area, *A*_hydro_(0).

|  |  | SA | CD2 | CD4 | CD45 |
| --- | --- | --- | --- | --- | --- |
| Theory<sup>a</sup> | $a$ [nm] | 2.8 | 1.5 | 1.5 | 1.5 |
| | $h_c$ [nm] | 5 | 7.5 | 11 | >15 |
| | $A_{\text{hydro}}(0)$ [nm<sup>2</sup>] | 296 | 299 | 513 | $\geq 820$ |
| Experiment<sup>b</sup> | $A_{\text{hydro}}(0)$ [nm<sup>2</sup>] | 300±29 | 324±18 | 505±40 | 1536±352 |
| | $h_{\text{Ahydro}}$ [nm]<sup>c</sup> | 5.1±0.4 | 8.0±0.3 | 10.9±0.6 | 22.2±3.1 |
| | $a_{\text{hs}}$ [nm]<sup>d</sup> | 3.2±0.4 | 5.3±0.2 | 3.7±0.3 | 8.0±0.3<sup>e</sup> |

<sup>a</sup> Theory from: Darst et al. (SA),<sup>4</sup> Davis and van der Merwe (CD2),<sup>16</sup> Yin et al. (CD4),<sup>8</sup> and Chang et al. (CD45).<sup>7</sup>
<sup>b</sup> Experimental values are given as mean ± SD.
<sup>c</sup> Estimated height value (mean ± SD) calculated from Eqs. S2 and S3 using the experimental $A_{\text{hydro}}(0)$ and the theoretical $a$ .
<sup>d</sup> Fit of $\mu^{\text{ex}}$ to a hard disk model, with $a_{\text{hs}}$ being the disk radius.
<sup>e</sup> Value at $c = 500$ molecules/ $\mu\text{m}^2$ . $a_{\text{hs}} = 5.3 \pm 0.1$ nm at $c = 6000$ molecules/ $\mu\text{m}^2$ .

### The interaction between rod-like molecules - CD2, CD4 and CD45

The interaction curves in Figure 3A level off at different concentrations, indicating that the chemical potential of the proteins will be different. This can also be observed in Figure 3B which shows the excess chemical potential for CD2, CD4 and CD45. The excess chemical potential for all three proteins increases considerably faster than it would if they were hard disks with a radius of 1.5 nm. CD2 and CD4 could be modelled as hard disks with a radius of 5.3 nm and 3.7 nm, whereas CD45’s excess chemical potential curve altogether deviates from that of a hard disk. The reason for this is protein glycosylation and flexibility.

### Protein glycosylation

An explanation for the larger effective radius for CD2 and CD4 can be due to protruding Nglycans from the protein core. The length of the sugars on CD2 and CD4 are of the order of 3 to 4 nm,^15^ and can thus significantly extend the lateral dimensions of the proteins. The larger radius of CD2 compared to CD4 would then be in agreement with CD2 being more glycosylated (four glycosylation sites for CD2 compared to two for CD4).^15^ To investigate this further, MC simulations were performed for a model protein, mimicking different amounts of glycosylation (Figure S5). Two parameters, the *projected area* and the *hard-disk area*, were introduced to characterize the interactions. The former is the total cross-sectional area of the protein and sugars, whereas the latter is the corresponding area of a hard disk with equal chemical potential at 1 *k*_B_*T*. The hard-disk area was found to be equal to the projected area at low sugar densities (Figure 3C). However, the hard-disk area can be significantly larger than the projected area at higher densities and approach the projected area of a disk with a radius equal to the protein radius + the sugar length; Figure 3C. The effect of glycosylation is significantly less pronounced if the protein can rotate freely around its attachment point, resulting in tilt angles between ±90°. The hard-disk area now depends on the volume of the sugar groups instead of their projected area (Figure S6). This is valid for the simplified systems modelled here were all the sugars are assumed to be in one plane of the molecule.

### Protein flexibility

Thermal motion will, if the molecule or attachment point at the surface is flexible, cause the molecule to rotate, or tilt, around its attachment point. This affects the effective width of the molecule on the surface. Figure 3D shows the results from different MC simulations where a rod-like molecule can rotate around its attachment point. For a rod with a 3:1 height-to-width ratio the effective width of the protein increases by 1.4, 1.7 and 2.3 times that of a rigid rod (at 1 *k*_B_*T*), when the proteins can rotate ±10°, ±50° and ±90° around their attachment point, respectively. The effective width increment also depends on the rod length, with longer rods showing a larger increase (Figure S7). The chemical potential curves for the latter differ from the hard-disk curves, adopting a near linear dependence on surface coverage (Figure 3D), indicating that the repulsive interactions increase constantly with surface coverage.

### The interaction between rod-like molecules - CD2, CD4 and CD45 (continued)

If CD2 and CD4 can rotate ±10° around the vertical axis as previously suggested by Polley et al.,^17^ and the effective area per sugar group is 4 nm^2^,^15^ this yields an effective hard-disk radius of 3.6 nm for CD4 and 4.9 nm for CD2 (1.4× the hard disk radius from Figure 3C). It was here assumed that the sugars on CD4 and CD2 are similar in size but that CD4 has two sugars, whereas CD2 has four.^15^ These values are in qualitative agreement with the experimental observations, illustrating how glycosylation can significantly increase the intermolecular repulsion between proteins.

The excess chemical potential for CD45 is considerably higher than for the other proteins, indicating that CD45 is more repulsive than CD2 and CD4. However, compared to a hard-disk model the relative repulsion (effective cross-sectional area) decreases at higher concentrations. This agrees with a model where CD45 can freely rotate around its attachment point at the surface (Figure 3D). The experimental data were fit to MC simulated curves of freely rotating rods of varying height-to-width ratio (Figure S7). The fit for a 40 nm long molecule gave a radius of 3.8 nm, similar to the effective radius of CD4. The effective cross-sectional area at low coverage (500 molecules/μm^2^) corresponds to 200 nm^2^ but is less than half (88 nm^2^) of this value at 6000 molecules/μm^2^. In fact, the effective cross-sectional area at 6000 molecules/μm^2^ is similar to the area taken up by CD2.

### Trapping two types of proteins

Experiments were also performed for SLBs containing CD45 together with the protein CD2 to investigate the distribution of two differently-sized proteins in the trap (Movie S3). Figure 4 shows the distribution of both CD2 (red) and CD45 (green) at various times after the trap is turned on. Both proteins are initially accumulating in the centre of the trap, however, CD2 are partially excluded from the centre as the concentration of CD45 increases. The observed ring-like distribution of CD2 around CD45 can be explained by a combination of (i) shielding of CD2 by the larger CD45 molecules and (ii) competition of the free area in the centre of the trap (see “*Double trap theory*” in the SI and Figure S8). The accumulation of CD45 also weakens (Figure S9) with an *A*_hydro_(0) value of approximately 1000 molecules/μm^2^, compared to 1500 molecules/μm^2^ without CD2. This agrees with CD2 partially shielding CD45 from the flow. Similar, but less pronounced, data could be observed when investigating the trapping of CD4 together with CD45 (Figure S10).

**Figure 4.**
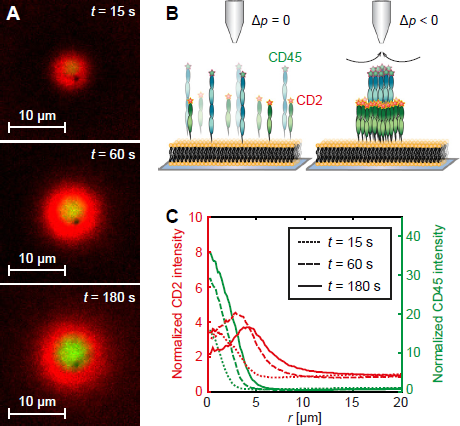
Simultaneous trapping of CD2 and CD45. **A)** Trapping of CD2 (red) and CD45 (green) after pressure is applied at *t* = 0. **B)** Scheme showing the double trapping. **C)** Radial line profiles of CD2 (red lines) and CD45 (green lines) at different times.

## Discussion

The hydrodynamic-trapping method allows us to investigate how membrane-anchored molecules interact over a wide range of concentrations, approaching concentrations of densely packed proteins. Such high surface coverages have previously been hard to study since they have required unpractically high concentrations of protein in solution when binding the molecules. In the current experiments the amount of immune-cell proteins used for each experiment were 10 to 100 ng, and the surface coverage in the centre of the trap exceeded 30%. Higher surface coverage can be obtained by increasing the applied pressure over the pipette, which in the current experiments was of the order of 10 kPa. Doubling the applied pressure for the SA experiments presented in Figure 2 would for example increase the maximum coverage in the trap from 32% to 48%, if the proteins still behave as hard disks with a radius of 3.2 nm. It is at these high coverages possible to investigate the aggregation of protein molecules and which conditions that favours this. For the investigated proteins it was only SA that could be aggregated, and this in turn required the use of high salt and low pH (650 mM NaCl and a pH of 3.6) and that the proteins were photobleached in the trap (Figure S1). The molecular reason for this behaviour remains to be investigated but demonstrates how the hydrodynamic trap can induce protein clustering under the appropriate conditions. Neither CD4 nor CD45, which have previously been argued to be able to form dimers,^18,19^ showed any trend to do so in our experiments. This indicates that if the molecules form dimers on the cell surface this is not driven by interactions between the extracellular parts of the molecules.

It has also been observed that deglycosylation of these molecules aids in forming protein crystals,^7,16^ thus indicating that the N-linked glycans protruding from the protein core increases the intermolecular repulsion. This was also confirmed by our experiments which combined with MC simulations allowed us to investigate how different degrees of glycosylation influences the interaction. It was found that even sugars constituting a minor part of the total protein weight can have a dominating effect on the intermolecular interaction. This effect is especially true for rigid molecules whose orientation relative to the lipid bilayer remains essentially constant. The effective repulsion of the glycosylated protein can here be calculated from the projected area of the sugar groups and will have a maximum relative effect for intermediate levels of glycosylation (Figure 3C). The effect of glycosylation will be less for flexible or freely rotating proteins, where in the latter case the increase in repulsion depends on the added total volume of the sugars. The difference comes from having all sugars in the same plane(s) with no or little rotation, but in different, non-coincident, planes when the molecule can rotate ±90° around its attachment point. Thus, glycosylation is expected to have the largest relative effect for rigid proteins.

Protein rotation, or flexibility, around the anchoring point at the surface can also on its own significantly affect the interaction potential and how it varies with protein coverage. The larger the rotation the higher is the interaction potential as shown in Figure 3D. This effect increases with the length of the studied protein. The reason for the extra repulsion is that the rotation causes the protein to take up a larger area on the lipid bilayer compared to when the protein is standing upright. This effect increases with the length of the protein and with the rotation angle from the lipid bilayer normal. When the concentration of proteins gets higher the initially non-constrained angles decrease resulting in the protein standing upright due to steric hindrance (maximum entropy). This transition costs in free energy and the interaction potential also has a different, more linear appearance vs protein coverage compared to the behaviour of the interaction potential for hard disks. This was also the appearance of the interaction potential obtained for the long CD45 molecule.

Both the orientation and the effective height of CD45 on the cell surface is presently unknown^7^ but has important consequences since the size-induced exclusion of these molecules from the contact between immune cells is the key mechanism in the kinetic-segregation model explaining T-cell triggering.^11^ Both the lower value of *A*_hydro_(0) compared to the predicted value for an upright molecule using Eq. 7 as well as the linear shape of the interaction potential vs protein coverage would be in agreement with a model where CD45, or its mucin-like region, is free to rotate around the attachment point, at least at low protein coverage. If the protein would dimerise or aggregate this would limit the possible angles that CD45 could rotate in thus increasing the free energy of the system. Protein flexibility will thus counteract aggregation in this case. However, when the protein coverage increases this will force CD45 to adopt a more upright position. The high concentration of CD45 and CD43 on the T-cell surface, approximately 1000 molecules/μm^2^ for each protein (Hui and Vale^20^ and S.J. Davis, personal communication), would result in the effective cross-sectional area of CD45 being approximately 40% lower than at infinite dilution. This indicates a reduction in allowed tilt angles and a more upright position of CD45 on the T-cell surface compared to on an SLB at low coverage.

We finally investigated how having more than one type of molecule on the surface affects the distribution in the trap. It was observed that the concentration of the largest molecule looked similar as when trapping the protein alone, while the smaller molecule got segregated from the centre of the trap (Figure 4). It is thus possible to locally segregate molecules based on size, with the largest molecules being in the centre of the trap. Since the effective size of a certain protein molecule can be increased by, for example, binding an antibody to the protein, coupled to a bead to increase its size further, this would make it possible to locally increase the concentration of a certain protein with respect to other molecules even in a more complex setting such as that on a live cell surface. The presented method thus opens for a deeper understanding of various processes revolving about how membrane-associated molecules interact. We also foresee how it can be extended to study a range of other interactions including, for example, dimer formation and aggregation of membrane-bound proteins.

## Methods

### Lipid and vesicle preparation

Vesicle solutions containing (i) 0.05% of 1,2-dipalmitoyl-sn-glycero-3-phosphoethanolamineN-(cap biotinyl) (sodium salt) (biotin-PE, Avanti^®^ Polar Lipids, Inc) mixed with 99.95% of 1-palmitoyl-2-oleoyl-sn-glycero-3-phosphocholine (POPC, Avanti^®^ Polar Lipids, Inc) or (ii) 5% or 10% 1,2-dioleoyl-sn-glycero-3-[(N-(5-amino-1-carboxypentyl)iminodiacetic acid)succinyl] (nickel salt) (DGS-NTA, Avanti^®^ Polar Lipids, Inc) mixed with 95% or 90% POPC, respectively, were prepared at a concentration of 0.5 mg/mL by the following protocol. The different lipids were first mixed in 100 µL chloroform. To remove the chloroform, the lipids were dried using a N2 gas flow for 10 minutes. After evaporation of chloroform the vesicles were suspended and thoroughly mixed in 1 mL of filtered (0.2 µm Minisart^®^ Syringe filter, Sartorius) washing buffer: 150 mM NaCl, 10 mM 2-[4-(2-hydroxyethyl)piperazin-1-yl]ethanesulfonic acid (HEPES, Sigma), pH 7.4 for all lipids. The vesicles were then incubated on ice for 1-2 h followed by tip sonification with a CV18 model tip sonicator (Chemical instruments AB) for 15 minutes with a pulse time of 10 s and an amplitude of 55%. The lipid stock solutions were stored at −20°C in chloroform and the vesicle solutions at 4°C until used.

### Cover slide preparation and protein loading

0.15 mm thick, round glass slides (number one coverslips Ø 25 mm, Thermo Fisher Scientific) were cleaned for 30 min in 80°C heated piranha solution (mixture of 75% sulfuric acid (99.9%, Sigma) and 25% hydrogen peroxide (30%, Sigma)). After the piranha wash the glass slides were rinsed for one minute on each side with deionised water and then dried with N2 gas. A silicon well (Silicon isolators, 12×4.5 mm diameter, 1.7 mm depth; Grace Biolabs) was cleaned with ethanol, paper dried and dust particles were removed with tape. The clean glass slide was pressed on the silicon well and placed in an Attofluor^®^ Cell Chamber (Thermo Fisher Scientific). The well was filled with washing buffer and left for sample preparation covered with a Petri dish to avoid dust in the buffer. Vesicles were diluted 1:10 in 30 µL of the washing buffer, added to the prepared well and incubated for one hour at room temperature. This allowed for formation of a fluid and continuous SLB on the glass surface. Non-ruptured vesicles were after the incubation washed away and the proteins were added to the well. 40 µL of Streptavidin, Alexa Fluor^®^ 647 conjugate (Thermo Fisher Scientific) were added at a concentration of 20 ng/µL and bound to the biotin-PE lipid bilayers.

Binding of proteins to the DGS-NTA lipids required a histidine tag covalently bound to the proteins. Thus, recombinant rat CD2 with a modified C-terminus containing a double histidine tag, as well as human CD45RABC and human CD4 containing one histidine tag, were made to bind to the DGS-NTA lipids. The proteins were fluorescently labelled with the dyes Alexa Fluor^®^ 488 and Alexa Fluor^®^ 647 using an Alexa Fluor antibody labelling kit from Thermo Fisher Scientific. All proteins were diluted in 150 mM NaCl and 10 mM HEPES, pH 7.4. Whereas 30 µL of 0.5 ng/µL CD2 or CD4 were loaded on the DGS-NTA lipid bilayer, 30 µL of 3 ng/µL of CD45 were used. The proteins were allowed to settle for one hour and excess proteins were washed away with washing buffer.

### Microscopy setup

The fluorescently-labelled molecules were studied with a Nikon Apo TIRF 60x magnification oil immersion objective on a Nikon Eclipse TE2000-U microscope equipped with a Hamamatsu ORCA-Flash4.0 LT Digital CMOS camera (C1140-42U). The sample was illuminated using Cobolt MLD compact diode lasers operating at a wavelength of 488 nm (20 mW) and 638 nm (140 mW). The acquired images were 512 pixels by 512 pixels with a pixel size of 0.22 µm by 0.22 µm.

For fluorescence recovery after photobleaching (FRAP) analysis a series of pre-bleached images was taken under the above stated settings. The field diaphragm was closed to a small circular area and left for six seconds to illuminate and bleach a small region on the visible SLB. In this time the ND filter (ND 2.0) was removed for three seconds from the beam path resulting in efficient bleaching of the illuminated area. The ND filter and field diaphragm were returned to their original positions and the recovery was followed for 106 seconds. For all experiments the time between frames was two seconds.

### Pipette pulling and characterization

Micropipettes with a size of 2-5 µm in inner tip diameter were pulled with a flaming/brown type micropipette puller (model P-97; Sutter Instruments) from borosilicate glass capillaries with an inner diameter of 0.58 mm and an outer diameter of 1.0 mm. An image of the pipette was acquired in bright field mode with a 20x magnification air objective (1.54 pixels/µm). The outline of the pipette was manually determined and can be described as multiple connected conical segments (Figure 5). The dimensions of the segments were used to determine the flow profile outside the pipette using finite element simulations (see “*Methods - Simulations of the flow and ion current in the pipette*”).

**Figure 5.**
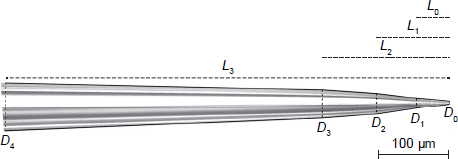
Image of a micropipette captured with a 20× magnification air objective. Solid lines denote the outer walls of the pipette, dashed lines denote the start of a new conical segment and the dotted lines represent the distance between the pipette tip and the conical segments.

### Protein accumulation under hydrodynamic forces

The pulled micropipette was mounted on a custom-made micropipette holder for the hydrodynamic trapping experiments. The holder can be moved in the *x*-, *y*- and *z*-direction with nm-precision using a 3D piezoelectric positioning system (P-611.3s NanoCube^®^XYZ-system, Physik Instrumente). Manual stages were used for more coarse positioning in the *x*- and *y-*direction and for coarse motion in *z*-direction a motorized stage from Thorlabs (T-cube Brushless DC Servo controller, PT1/M) were used. This assures that the mounted pipette can be centred over the sample. In addition, an Ag/AgCl electrode was inserted in the pipette. To measure the distance of the pipette to the SLB the ion current between the electrode in the pipette and an Ag/AgCl reference electrode in the bath solution around the pipette was measured. The electrode in the micropipette was chlorinated before each experiment using bleach to guarantee a stable ion current between the two electrodes. When the pipette approaches the surface, the resistance between the two electrodes increases. This increase, around 0.5 - 1% in the current experiments, can be converted to an actual distance of the pipette to the surface when the dimensions of the pipette are known.^2^ The pipette was then moved away between 2 µm to 6 µm to create a bigger area for the trapping region.

Accumulation of proteins was initiated by applying negative pressure over the micropipette. The proteins were accumulated under different pressures: −3±0.3 kPa, −9.7±0.3 kPa and −19.4±0.3 kPa. Between each pressure change, and measurement, the concentration of the proteins in the trap had reached equilibrium. Finally, the proteins were released by turning off the pressure. The accumulation was captured with the microscope settings stated above, but with a time frame of 15 seconds between each image.

### Analysis of the trapping data

The acquired fluorescence images were analysed using a custom-written MATLAB^®^ (MathWorks Inc.) program to give an interaction curve for each trapping event. A fluorescence image of the trapped region was acquired after the concentration of proteins had reached steady state and the intensity was radially averaged around the centre of the trapped region. The radial fluorescence intensity was next converted to a molecular density by a conversion factor obtained from single molecule imaging as described in the section “*Methods - Conversion between fluorescence intensity and protein density*”. Corresponding values for *ε*_hydro_ at different radial positions were determined as described in the section “*Methods - Determining *ε*_hydro_ and *σ*_hydro_ from finite element simulations*”. Plotting the molecular concentration *c* as a function of *ε*_hydro_ finally gave the interaction curve.

### Conversion between fluorescence intensity and protein density

A 1 pM solution of labelled proteins was added to a silicone well on a glass cover slide. The proteins were allowed to bind for 10 minutes after which the protein solution was replaced with buffer solution. This resulted in 100-200 protein molecules binding randomly in the microscope field of view, where each molecule could be individually distinguished. At least three different areas were analysed for each protein. The molecules were detected and counted using an implementation of the particle detection algorithm by Crocker and Grier.^21^ The sum of the intensity from each molecule, within a radius of 4 pixels from the centre of each detected molecule, was determined and subtracted with the average intensity outside the molecules to correct for non-zero background. The fluorescence intensity per pixel, *I*, was converted to a protein density, *c*, using:

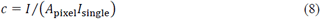

where *A*pixel is the area of a pixel in the image and *I*single is the average intensity from a protein molecule. The standard deviation of the obtained *I*single values, from the different experiments, was 5% or smaller. The obtained conversion factors were finally corrected for all protein molecules not being labelled. This was mainly necessary for CD2 where the average number of dyes per molecule was 0.75. In short, the step change in intensity of the protein when subjected to photobleaching, *I*step, was determined which corresponds to the intensity from one dye. The mean number of dyes per protein was determined using a NanoDrop spectrophotometer (Thermo Fisher Scientific). Multiplying *I*step with the average number of dye molecules per protein gave the corrected value of *I*single for CD2.

### Determining *ε*_hydro_ and *σ*_hydro_ from finite element simulations

Different factors such as the geometry of the pipette, the distance between the pipette tip and the SLB and the pressure applied over the pipette affects *ε*_hydro_. The distance between the tip of the pipette and the underlying SLB can be related to the change in ion current as the pipette approaches the surface using finite element simulations.^2^ To avoid having to do a new simulation for each experiment, the following approximate expression, valid for the low-tapered pipettes used in this work, was used:

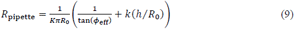

where *R*pipette is the electric resistance over the pipette, *K* is the ion conductivity in the medium, *R*0 the inner tip radius of the pipette and *k*(*h*/*R*0) a function of the dimensionless parameter *h*/*R*0 with *h* being the distance between the tip of the pipette and the SLB. The inner tip radius is obtained from *D*0 in Figure 5 by multiplying with 0.29, where it is assumed that the ratio between inner and outer diameter of the pipette is constant along the length of the pipette (*R*inner/*R*outer = 0.58). The parameter *ϕ*eff is the voltage inner half cone angle of the multi-segmented pipette, and is defined by:

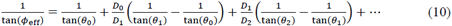

where

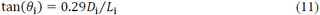

and it is assumed that the ratio between inner and outer diameter of the pipette is constant along the length of the pipette. It is enough to do a series of finite element simulations at different heights *h* above a surface for a reference pipette after which Eq. 9 can be used to determine *R*pipette and its dependence of *h* for an arbitrarily-sized pipette.

The value for *ε*_hydro_, or *σ*_hydro_, can also be determined from finite element simulations.^2^The shear force *σ*_hydro_ will, in the creeping flow regime, be given by:

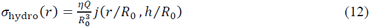

where *η* is the viscosity of the fluid and *j* is a function of the two parameters *r*/*R*0 and *h*/*R*0, where *r* is the radial distance from the centre of the trap at the surface and *h* is again the distance between the pipette tip and the surface. The parameter *Q* is the liquid flow rate out of the pipette, which for the small cone angles considered in this work can be shown to have the following approximate dependence on the applied pressure Δ*p* over the pipette:

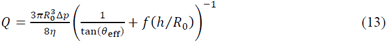

where *f* is a function that only depends on the parameter *h*/*R*0 and *θ*eff is the pressure inner half cone angle which for a pipette with multiple segments is given by the expression:

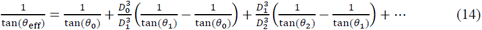

The value for *σ*_hydro_ as a function of *r* was determined for a reference pipette at a series of different values of *h* using finite element simulations as previously described.^2^ With this tabulated data for the reference pipette and Eq. 12 and Eq. 13 it is possible to determine *σ*_hydro_for an arbitrarily-sized pipette at different distances *h* from the surface.

### Metropolis-Hastings Monte Carlo simulations

All MC simulations were performed using the Faunus software package.^22^ Molecules were treated as a sequence of connected spheres, the first grafted to a hard, planar surface. Molecules were allowed to rotate and move along the surface during each MC step, and periodic boundary conditions were applied in the *x*- and *y*-direction. A purely repulsive, truncated and shifted Lennard-Jones pair potential was used for inter-sphere interactions.^23^ This resulted in a slightly larger disk (7.5% larger radius) than when using a hard disk potential, which was corrected for by multiplying all surface coverages with 1.075×1.075. Electrostatic interactions were neglected due to the high salt concentration used and small surface charge density of the proteins.

In order to constrain rotation, the sphere furthest from the surface may be exposed to a quadratic potential: *u* = *kf* (*z-Lmax*)^2^, where *z* is the distance from the surface, *Lmax* is the length when standing perpendicular to the surface, and *kf* is a spring constant. Thus, for the limit *kf* → 0 the molecules are freely rotating, while if *kf* →∞ they stand perpendicular to the surface. We investigated both soft (*kf =* 1 *kBT*/Å^2^), hard (*kf =* 5 *kBT*/Å^2^) and very hard (*kf =* 50 *kBT*/Å^2^) spring constants, giving rise to 95% probabilities of ±50°, ±10° and 0° degrees deviations from the surface normal, respectively. MC simulations with 1 million steps were run with different types of studied molecules, investigating the effect of length and topology of the molecule on the chemical potential of the system. Excess chemical potential curves as a function of surface coverage was calculated by the Widom insertion method.^23^ In the basic study, all spheres of the molecule had a diameter of 0.1 nm and the box size was 2×2×2 nm^3^. A bigger simulation box was used (4×4×4 nm^3^ or 6×6×6 nm^3^) when modelling the effect of glycosylation. The protein part of the molecule was here substituted by three connected spheres with a diameter of 0.3 nm while the sugars were approximated with two connected spheres with a diameter of 0.16 nm (Figure S5). The topology of the molecules was fixed during the simulation and no additional internal flexibility was allowed. The surface coverage, where each of the excess chemical potential curves was 1 *k*B*T*, was interpolated from the simulated curves in Figure S11 and the ±10° data was used to produce Figure 2C. The data was fit to the following empirical formula:

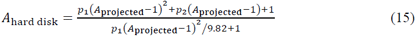

where *A*hard disk and *A*projected is the *hard-disk area* and *projected area*, respectively, normalized to the projected area without sugars. Equation 15 levels of at 9.82, which correspond to the (normalized) projected area of a sphere with the radius given by the protein radius + sugar length in our simulations (Figure S5), and increases approximately linearly with *A*projected at low levels of glycosylation. The coefficients *p*1 = 0.66 and *p*2 = 0.95 were determined by fitting Eq. 15 to the experimental data.

## Acknowledgements

This work was supported by grants from the Swedish Research Council (number: 621-2014-3907), the Royal Physiographic Society in Lund, the Crafoord Foundation, the Royal Swedish Academy of Sciences, and the Wellcome Trust.

## Author Contributions statement

VJ did all trapping experiments and analysed part of the data, JH and ML did the MC simulations, AMS and SJD made the immune-cell proteins, PJ did most of the data analysis and finite element simulations, PJ and VJ wrote the paper.

